# Gut microbes predominantly act as symbiotic partners rather than raw nutrients

**DOI:** 10.1101/2023.02.08.527620

**Authors:** Nuno Felipe da Silva Soares, Andrea Quagliariello, Seren Yigitturk, Maria Elena Martino

## Abstract

Animals and their gut microbes mutually benefit their health. In this frame, nutrition has a central role by directly affecting both host and microbes’ fitness and their effects. This makes nutritional symbioses a complex and dynamic tri-system of diet-microbiota-host. Despite recent discoveries on this field, full control over the interplay among these partners is challenging and hinders the resolution of fundamental questions, such as how to parse the gut microbes’ effect as raw nutrition or as symbiotic partners? To tackle this, we made use of the well-characterized *Drosophila melanogaster*/*Lactiplantibacillus plantarum* experimental model of nutritional symbiosis to generate a quantitative framework of gut microbes’ effect on the host. We show that the beneficial effect of *L. plantarum* strains primarily results from the active relationship as symbionts rather than raw nutrients, regardless of the bacterial strain. Metabolomic analysis of both active and inactive bacterial cells further demonstrated the crucial role of the production of beneficial bacterial metabolites, such as N-acetylated-amino-acids, as result of active bacterial growth and function. Altogether, our results provide a ranking and quantification of the main bacterial features contributing to sustain animal growth. We demonstrate that viability is the predominant and necessary variable involved in bacteria-mediated benefit, followed by strain-specific properties and the nutritional potential of the bacterial cells as direct energy source. This contributes to elucidate the role of beneficial bacteria and probiotics, creating a broad quantitative framework for host-gut microbiome that can be expanded to other model systems.

## Introduction

The complex relationship between animals and their microbiota is one of the most ubiquitous and relevant biological interactions observed in nature. It can range from neutral, mutually beneficial to harmful and pathogenic^1^. The gut microbiota largely influences animal’s health by enhancing host metabolism, growth and development, priming the immune system and protecting the host from invasion by pathogens^2–8^. The establishment and evolution of such relationships result from a combination of genetic and environmental factors. Specifically, host factors (i.e., developmental stage, genetic background, etc.) have been shown to influence gut microbiome composition and function across different animal species (e.g., human, mouse, zebrafish, chicken, cattle, and swine)^9–15^. At the same time, environmental factors (e.g., diet, co-habitation, abiotic factors, etc.) were found to best describe microbiome composition in humans and animals^16–20^. In this frame, the host diet has a major impact on gut microbiota diversity, evolution and activity^18,21–27^. Short-term dietary interventions have been shown to induce rapid, albeit transient, alterations on the gut microbiota diversity^27–29^. In addition, dietary composition can alter the gut microbes’ physiology and their ultimate effect on host health. The presence of certain microbial species in the animal gut microbiome is sufficient to rescue nutritional deficiencies by converting and providing essential nutrients to its host, such as amino-acids and vitamins^30–32^. Conversely, scenarios where microbiome composition and function are severely unbalanced (i.e., gut dysbiosis) have been associated to several diseases, including obesity^33,34^, chronic inflammatory diseases^35–38^, and metabolic disorders^21^. Therefore, the gut microbiota represents a fundamental intermediary between the host and its diet. Nevertheless, it is difficult to ascertain if given host phenotypes (e.g., development, aging) are attributable to bacterial-mediated compensation for nutritional deficiencies (i.e., bacteria serve as mere nutritional sources) or, instead, caused by direct bacteria-elicited responses.

Here, we employed a well-established model of host/microbe symbiosis, *Drosophila melanogaster* associated with *Lactiplantibacillus plantarum*, to define and quantify the role of gut microbes as beneficial symbionts.

Drosophila carries simple bacterial communities, mostly dominated by *Acetobacter* and *Lactobacillus* species^39^. It is widely recognized that gut microbes modulate numerous fly life-history traits, such as juvenile growth, lifespan, and even behavior^30,32,40,41^. This is mainly due to their ability to provide their host with access to essential nutrients, particularly in nutritionally challenging conditions^32–34,40–42^. *Lactiplantibacillus plantarum*, an ubiquitous bacterial species and one of the most common *D. melanogaster* gut symbionts, has been demonstrated to be able to promote animal growth in conditions of protein scarcity^40,43,44^. Importantly, such beneficial effect is strain-specific^44–48^. In this frame, we have previously shown that non-beneficial *L. plantarum* strains are able to improve their growth-promoting effect by adapting to the host nutritional environment^26^. The improved adaptive potential allows bacteria to reach higher bacterial loads and produce a higher concentration of beneficial metabolites, ultimately becoming more beneficial. Indeed, recent research has reported that microbes act primarily as a critical source of protein and that the beneficial effect of gut microbes relies on the bacterial ability to thrive in the fly nutritional environment^49^. Microbial abundance is thus critical for promoting fly growth and development^26,43,49^. At the same time, Drosophila gut microbes are able to actively influence the host response, by modulating Tor and insulin signaling pathways, promoting intestinal peptidase expression, boosting the immune response, as well as cooperating among them^31,40,41,45,46,50–53^. Altogether, these studies demonstrate that microbes possess a dual role in promoting host development. Yet, the extent to which gut microbes act as beneficial active partners and mere nutritional sources remains elusive.

Investigating the role of gut microbes as source of nutrients has been mainly performed by testing the effect of heat-killed bacteria on host traits (e.g., development, lifespan)^30,40,43,48,49,54^. However, heat treatments have profound effects on the structural properties of bacteria, significantly altering membranes, proteins and enzymes^55,56^. Such structural modifications may affect the nutritional value of bacterial cells, impairing the direct comparison of the effects between live and dead bacteria. To overcome this limitation, we established a quantitative framework of host/microbe relationship by associating *Drosophila* larvae with defined and stabilized live bacterial loads. We used erythromycin, a bacteriostatic antibiotic, to control and monitor both the bacterial physiological status and loads during host-microbe associations and test their effect on host development (i.e., larval growth and developmental time). Given the strain-specific effect of gut microbes, we analyzed two pairs of isogenic *L. plantarum* strains, each pair including one highly and one slightly beneficial strain in promoting *Drosophila* larval growth^26,45^. Our results show that bacteria viability (i.e., the capacity to actively interact with the host) is the predominant factor in microbe-mediated growth-promotion. This feature overcomes other crucial, albeit less important, microbial features, such as biomass (i.e., nutritional source) and strain-specific characteristics (i.e., cell features, ability to thrive in the environment). This study clarifies and ranks the multiple roles of gut microbes in promoting fly growth and establishes a new method to evaluate the effect of microbial biomass in host physiology, over the use of treated bacteria.

## Results

### Live and stable bacteria are unable to rescue larval size to standard mono-association levels

To investigate the nutritional value of bacterial cells, we sought to obtain unmodified bacterial cells whose growth was impaired (i.e., alive but stable). To this end, we first tested the effect of five bacteriostatic compounds (e.g., chloramphenicol, erythromycin, sulfamethoxazole, tetracycline and trimethoprim) on *L. plantarum* NC8 (*Lp*^NC8^)^57^ and *L. plantarum* NIZO2877 (*Lp*^NIZO2877^)^58^ growth using minimum inhibitory concentrations (MIC) tests. The MIC is defined as the lowest concentration of the antimicrobial agent where no bacterial growth is observed under defined conditions^59^. All tested antibiotics inhibited bacterial growth, except for sulfamethoxazole, to whom *L. plantarum* showed resistance. We then selected erythromycin as suitable bacteriostatic to control *L. plantarum* loads in association with Drosophila larvae, being the antibiotic with the lowest effective concentration (Supplementary Table 1).

To test whether bacterial cells remained alive and stable (i.e., non-dividing) in presence of erythromycin, we associated 10^7^ CFU/ml of *Lp*^NC8^ with low-protein fly food containing 20 μg/ml of erythromycin with and without Drosophila larvae, and monitored bacterial loads over the course of 6 days. As expected, we detected bacterial growth in absence of erythromycin, regardless of host presence (Supplementary Figure 1A). Specifically, at day 6, *Lp*^NC8^ showed 1-log increase in bacterial load in presence of Drosophila larvae compared to the experimental condition where the host was absent (Supplementary Figure 1A), further proving the known beneficial effect of the fruit fly towards bacterial proliferation^60^. When erythromycin was present, the bacterial load remained stable in absence of the larvae and it slightly decreased in presence of Drosophila (Supplementary Figure 1A). Following these results, we hypothesized two possible scenarios: 1) bacterial growth was impaired by erythromycin and most cells remained alive without dividing; 2) a bacterial sub-population died and it was replaced by newly divided live cells. The latter scenario would then create the same live bacterial load, with an increased total biomass. To test our hypotheses, we quantified the total bacterial DNA on low-protein diet in presence of erythromycin over the course of 3 days using qPCR. While we observed an increase in bacterial DNA concentration in absence of the bacteriostatic antibiotic, the total bacterial DNA level remained stable when erythromycin was present (Supplementary Figure 1B). Taken together, these results demonstrate that, in presence of erythromycin, *L. plantarum* cells remain alive and stable on low-protein fly food, regardless of the host presence.

We next investigated the effect of bacterial biomass and function on larval growth. To this end, we analysed three different bacterial states: 1) alive and growing (mono-association with live bacteria) to test for bacterial biomass and function, 2) alive and stable (mono-association with live bacteria in presence of erythromycin) and 3) dead and stable (mono-association with dead bacteria) to test for the effect of bacterial biomass. For the latter condition, we employed two different methods of bacterial cell killing (i.e., heat and UV treatment) to account for any potential bias derived from each method. The latter two conditions allowed to compare the effect of live, yet stable, biomass with dead biomass. We tested four different *L. plantarum* strains for each condition to account for bacterial strain-specific effect: *Lp*^NC8^ (strong growth-promoting strain), *Lp*^Δdltop^ (weak growth-promoting strain, isogenic to *Lp*^NC8^), *Lp*^NIZO2877^ (weak growth-promoting strain) and *Lp*^FlyG2.1.8^ (strong growth-promoting strain, isogenic to *Lp*^NIZO2877^)^26,45^. 10^7^ CFU/ml of each strain (live or dead) were associated with Drosophila embryos in all conditions and larval size was measured at day 4 and day 7 after mono-association. Our results show that Drosophila growth was maximized in presence of live and growing bacteria both 4 and 7 days after bacterial association (Figure 1, Supplementary Figure 2A). Contrarily, dead bacteria were not able to promote Drosophila growth, regardless of the killing method, as larvae associated with heat-killed or UV-treated bacteria showed the same length as GF larvae (Figure 1, Supplementary Table 2, Supplementary Figure 2A). Interestingly, in case of *Lp*^NC8^ and *Lp*^Δdltop^ associations, larvae associated with live and stable bacteria (i.e., growing on fly food supplemented with erythromycin) displayed a significant increase in length compared to larvae associated with dead bacteria, while no significant difference was detected in *Lp*^FlyG2.1.8^- and *Lp*^NIZO2877^-associated larvae (Figure 1). Such difference was visible 7 days after mono-association, but not at day 4 (Figure 1, Supplementary Figure 2A). Of note, the presence of the bacteriostatic did not show any significant effect on larval growth (Supplementary Figure 2B). Altogether, these results show that the increase in bacterial biomass is key to promote larval growth.

**Figure 1.**
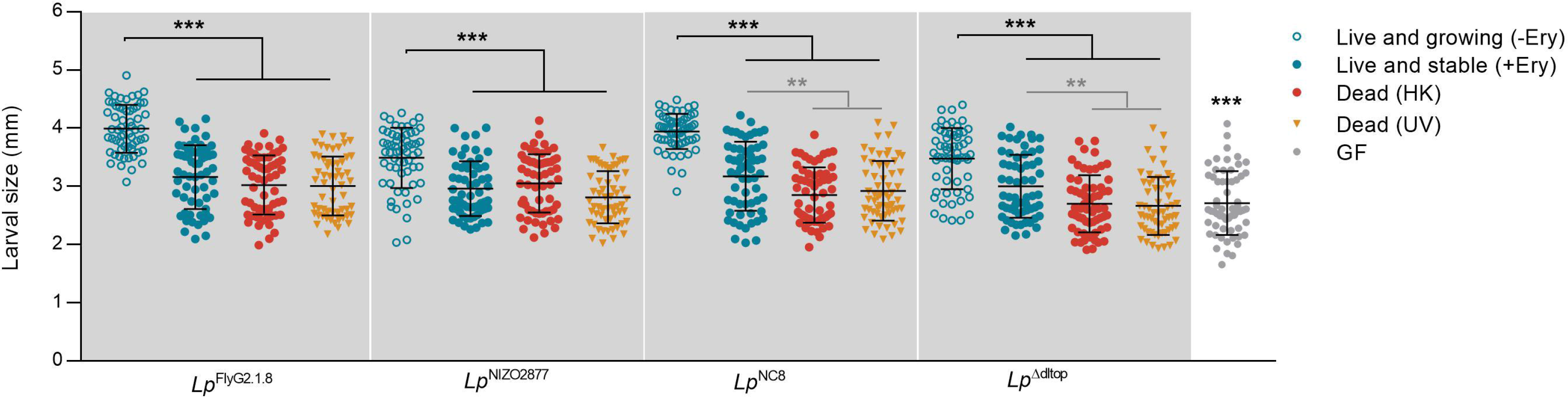
Larval longitudinal length measured 7 days after inoculation with 1) live bacterial strains on LPD without erythromycin (Live and growing −Ery); 2) live bacterial strains on LPD supplemented with 20μg/ml of erythromycin (Live and stable +Ery); 2) heat-killed bacterial strains on LPD without erythromycin (Dead – HK); 3) UV-treated bacterial strains on LPD without erythromycin (Dead – UV); and 4) PBS on LPD without erythromycin (GF condition). Asterisks illustrate statistically significant difference on pairwise intra-strain comparisons between standard mono-association (Live and growing -Ery) and the respective treatments (including GF) (***: p<0,0001, **: p<0,005). Center values in the graph represent means and error bars represent SD.

### Live and growing bacteria are necessary to promote larval growth

We next asked to what extent the growth-promoting effect provided by increasing bacterial loads relied on *i)* the active interplay between live bacteria and the host (i.e., production of metabolites, stimulation of host response, etc., resulting from live and dividing bacteria) and *ii)* the mere increase in bacterial biomass, regardless of bacterial vitality. To this end, we added 1) live and non-dividing bacterial cells and 2) dead bacterial cells on fly food containing Drosophila embryos (with and without erythromycin to test for any effect due to the presence of the antibiotic) every 24 hours for 7 days. The first condition allowed us to recapitulate standard bacterial growth, while the second condition allowed to analyze only the effect of the increase in bacterial biomass. The number of bacterial cells inoculated in each treatment was calculated considering the different growth dynamics of the four *L. plantarum* strains on LPD (Supplementary Figure 3A) in order to reach the same loads of a standard mono-association. No significant differences in bacterial CFUs were detected between standard mono-association and recurrent supplementation of live bacteria (Supplementary Figure 3B). The recurrent addition of live and stable bacteria provided the same effect on larval growth as standard mono-associations for all strains tested at both times of development (Figure 2A, Supplementary Figure 4). Consistent with previous studies, we found a significant correlation between larval size and bacterial counts, with strains reaching higher loads being more beneficial (*Lp*^FlyG2.1.8^>*Lp*^NC8^>*Lp*^NIZO2877^> *Lp*^Δdltop^, Figure 2B)^43,49^. Notably, the daily addition of inert bacterial biomass did not recapitulate the growth-promoting effect provided by live bacteria (Figure 2A, Supplementary Figure 4). In details, in case of *Lp*^FlyG2.1.8^- and *Lp*^NIZO2877^-associations, the addition of dead bacteria did not confer any beneficial effect, as larvae associated with the respective strain showed the same length as GF larvae. Instead, a slight increase in larval length was detected in case of daily addition of HK *Lp*^NC8^ and *Lp*^Δdltop^ strains. This suggests that the effect of bacterial biomass on larval growth is strain specific, with some bacterial cells (e.g., *Lp*^NC8^ and *Lp*^Δdltop^) providing more nutrients *per se* than others (e.g., *Lp*^FlyG2.1.8^ and *Lp*^NIZO2877^). Nevertheless, the artificial increase of dead cells to the equivalent amount of live cells was never sufficient to recapitulate the effect of live cells on larval growth, regardless of the bacterial strain.

**Figure 2.**
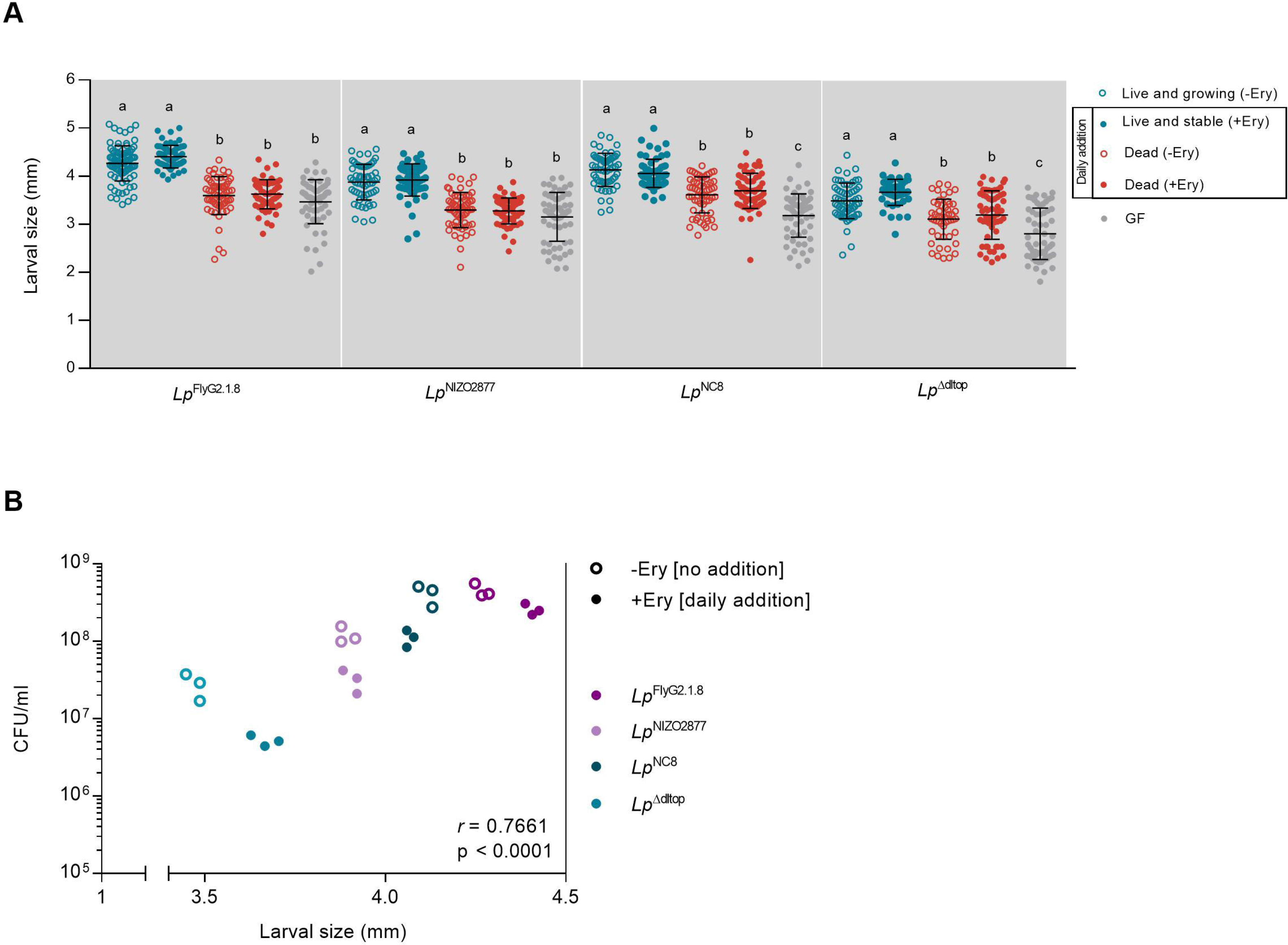
(A) Larval longitudinal length measured 7 days after inoculation with 1) live bacterial strains on LPD without erythromycin (Live and growing -Ery); 2) daily addition of live bacterial strains on LPD supplemented with 20μg/ml of erythromycin (Live and stable +Ery); 2) daily addition of heat-killed bacterial strains on LPD without erythromycin (Dead -Ery); 3) daily addition of heat-killed bacterial strains on LPD supplemented with 20μg/ml of erythromycin (Dead +Ery) and 4) PBS on LPD without erythromycin (GF condition). Different letters indicate statistically significant differences at p < 0.05. Center values in the graph represent means and error bars represent SD. (B) Microbial population size (CFU/ml) compared with average larval size from fly food containing the larval population showed in A. Microbe counts and larval size have been measured on day 7 after mono-association. -Ery, standard mono-association on LPD without daily addition of *Lp*. +Ery, mono-association on supplemented with 20μg/ml of erythromycin and daily addition of the respective *Lp* strain to recapitulate standard *Lp* growth (*r*=0.7661, p>0.0001, Spearman’s correlation efficient).

### Viability is the primary feature explaining bacteria-mediated benefit

Altogether, our results demonstrate that the increase in live bacterial biomass is fundamental to promote Drosophila growth. In addition, we further show that the growth-promoting effect mediated by bacteria is strain-specific and it likely relies on intrinsic features of the bacterial cells (Figure 2A). With the aim of identifying the most important predictors of bacteria growth-promoting ability, we measured the relative importance of each bacterial variable (i.e., biomass, viability) using Random forest (RF) classification^61^ at day 4 and 7 after bacterial association. All bacterial features experimentally tested in this study have been considered in the RF analysis: 1) viability, 2) biomass, 3) strain specificity and 4) antibiotic presence (Table 1).

**Table 1.**
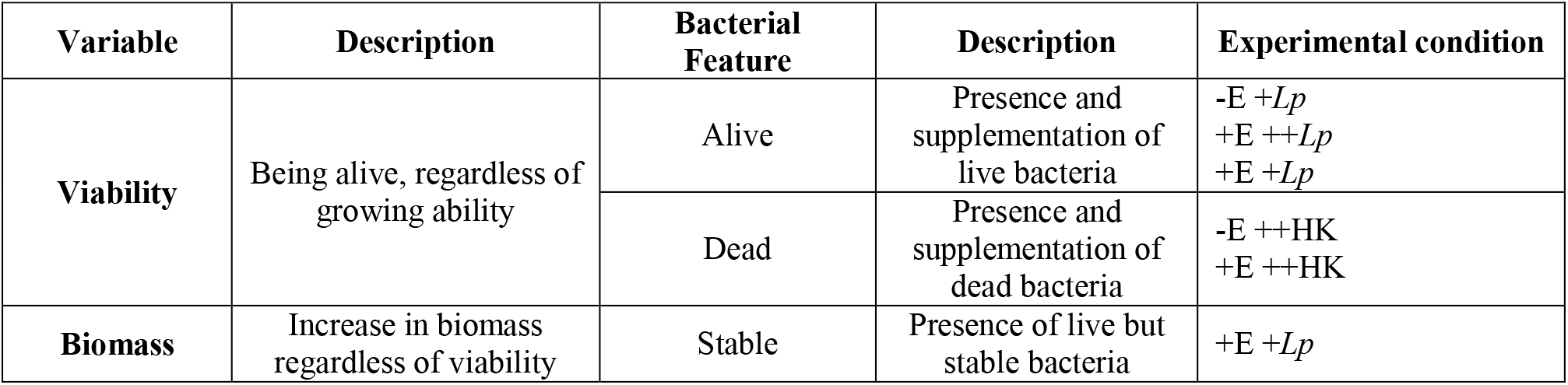
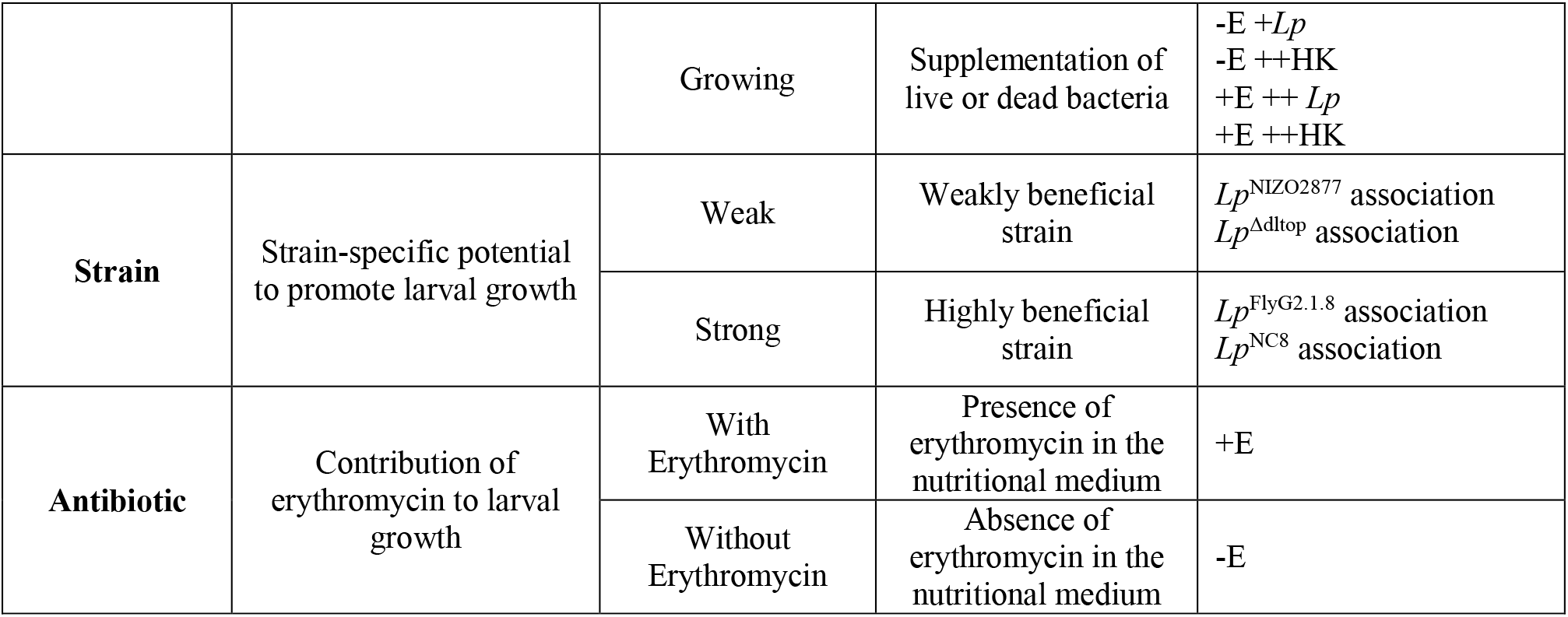
Description of the bacterial features tested for their relative importance in contributing to larval growth. −E: Absence of erythromycin in the nutritional medium; +E: presence of erythromycin in the nutritional medium; +*Lp*: single inoculation of *L. plantarum* at day 0 of mono-association; ++*Lp*: daily addition of *L. plantarum*; ++HK: daily addition of heat-killed bacterial cells.

Bacterial viability was ranked as the most important variable in promoting larval growth at both time points (Figure 3A, B, C), regardless of the bacterial strain (Supplementary Figure 5), contributing to 64% and 54% of larval development at day 4 and 7, respectively (Figure 3B). Instead, strain-specific features, as well as the mere increase in bacterial biomass (i.e., nutritional potential of bacterial cells), were attributed a significantly lower rank (Figure 3 A,B,D,E). Specifically, although the increase in bacterial biomass significantly promotes larval growth (Figure 3E), its contribution in determining the phenotype is remarkably lower than bacterial viability (Figure 3A,B). In addition, at early phases of larval growth (i.e., 4 days after bacterial association), bacterial biomass has been ranked as the least important feature (Figure 3A,B). Finally, the presence of erythromycin was found not to significantly alter larval growth (Figure 3A, E).

**Figure 3.**
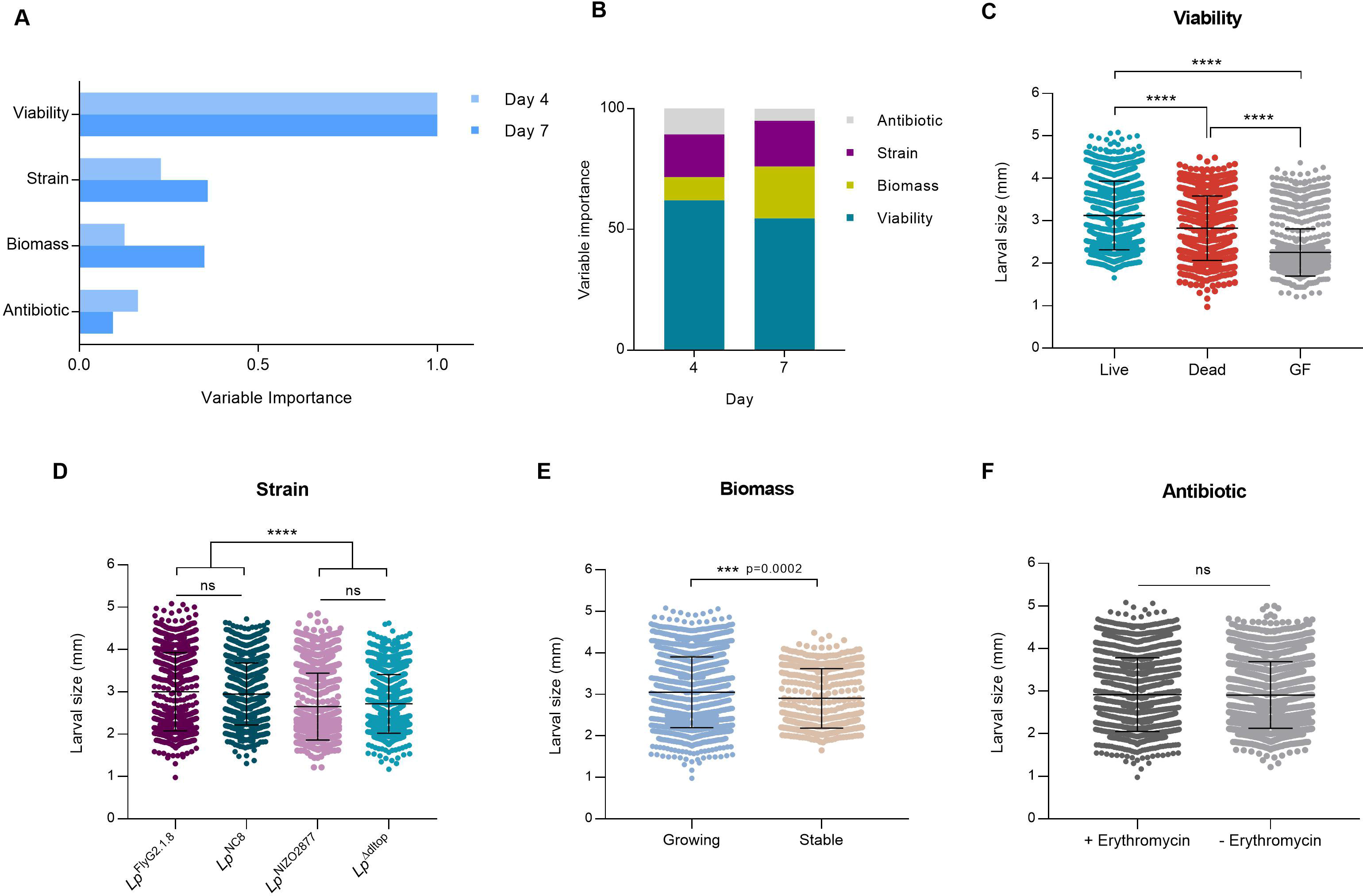
(A) Variable importance obtained through random forest analysis for each bacterial feature in promoting larval growth at day 4 and 7 after bacterial association. (B) Random Forest variable importance results as obtained from *ranger* function in R (*ranger* package). Variable importance has been scaled and transformed in percentage to describe percentage contribution of variable to the larval size. (C,D,E,F) Larval longitudinal length measured 7 days after inoculation with: (B) live bacterial strains (experimental conditions: -Ery [standard mono-association], +Ery [no daily addition], +Ery [daily addition]), dead bacterial strains (-Ery [daily addition of HT bacteria], + Ery [daily addition of HT bacteria]), and PBS (germ-free GF conditions); (C) the four *L. plantarum* strains, each including all experimental conditions tested (-Ery [standard mono-association], +Ery [no daily addition], +Ery [daily addition], -Ery [daily addition of HT bacteria], + Ery [daily addition of HT bacteria]); (D) increasing bacterial biomass (growing, experimental conditions: -Ery [standard mono-association], +Ery [daily addition], -Ery [daily addition of HT bacteria], + Ery [daily addition of HT bacteria]) and stable bacterial biomass (+Ery [no daily addition], -Ery [single inoculation of HT bacteria]); (E) live or dead bacterial strains on LPD with (+ Erythromycin) and without erythromycin (-Erythromycin). Asterisks illustrate statistically significant difference on pairwise intra-strain comparisons (****: p<0,0001, ns: not significant). Center values in the graph represent means and error bars represent SD.

### Live and growing bacteria produce more beneficial metabolites

We next investigated which molecules exclusively produced by live and growing bacteria would be responsible for improved larval growth. To this end, we analyzed the metabolome of Drosophila diets colonized with either *Lp*^NIZO2877^ or *Lp*^FlyG2_1.8^ strains, with and without erythromycin. As expected, the metabolome profiles of the two strains in absence of the antibiotic can be clearly distinguished from those in stable conditions (i.e., presence of erythromycin) (Figure 4A, Supplementary Table 3). We observed a significant increase in the levels of lactic acid, 3-phenyllactic acid and N8-acetylspermidine in presence of live and growing bacteria, while amino-acids (e.g., methionine sulfoxide and aspartic acid), organic acids (e.g., citric acid, malic acid, pyruvic acid) and sugars (e.g., hexose dimer) were significantly depleted (Figure 4B). Such compounds were mainly derived from the diet, as their concentration was significantly different between samples containing live bacteria and the diet alone (Figure 4C, Supplementary Table 3). The trend in metabolite production appears to be overall comparable between strains in the respective condition (e.g., *Lp*^FlyG2_1.8^ -E *vs Lp*^NIZO2877^ −E; Figure 4A, B). Nevertheless, we detected some inter-strain differences. The strong growth-promoting strain *Lp*^FlyG2_1.8^ was able to produce a significantly higher array of beneficial molecules, compared to *Lp*^NIZO2877^ (Figure 4B,C, Supplementary Table 3), while *Lp*^NIZO2877^ appears to consume more nutrients from the diet, such as amino acids, sugars, vitamins and nucleotides (Figure 4B,C, Supplementary Table 3).

**Figure 4.**
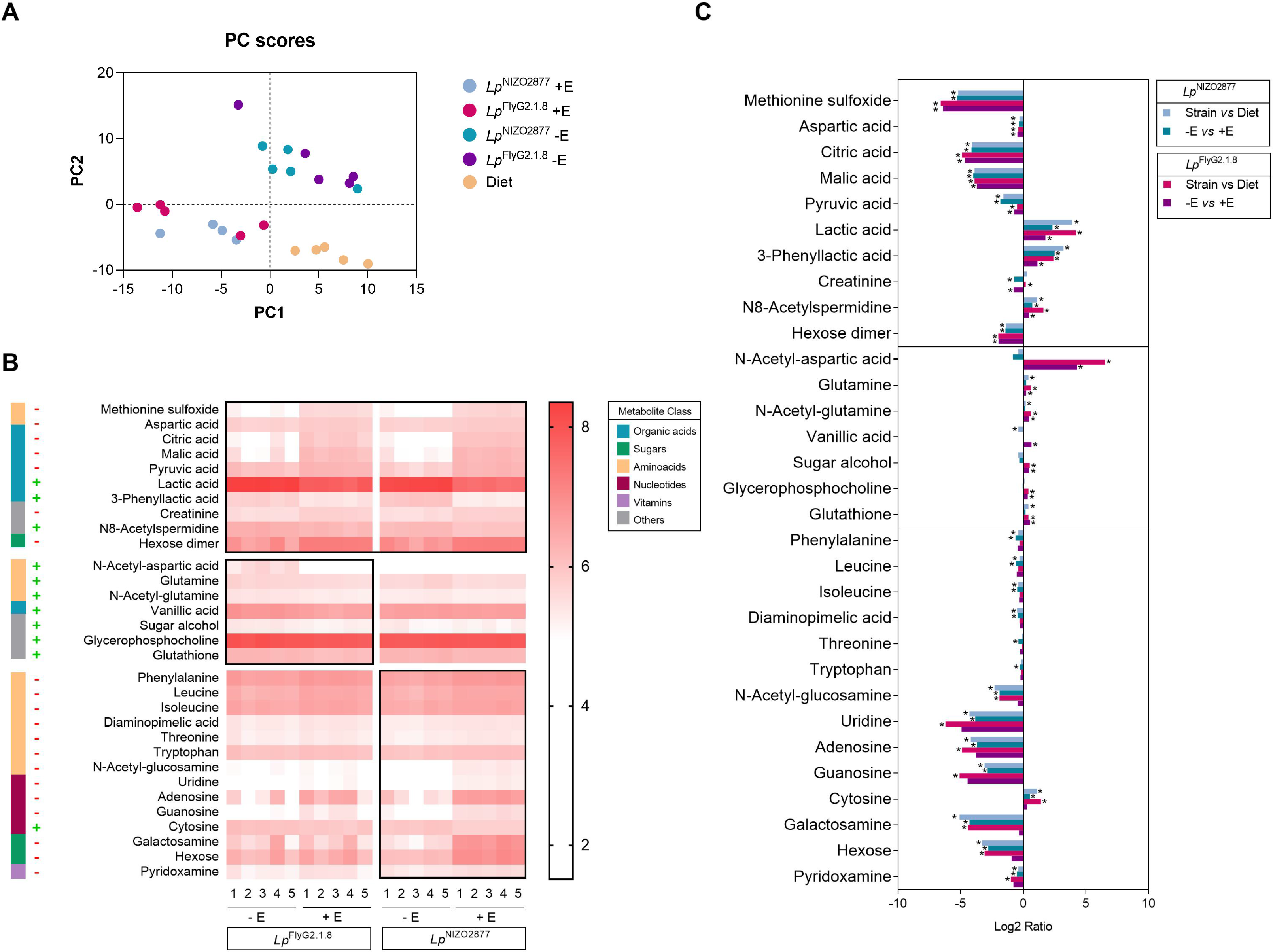
Metabolic differences between active and stable bacteria. (A) Principal Component Analysis (PCA) of the 5 experimental conditions based on the metabolic profiles. As expected, the presence of antibiotic discriminates samples on both axes (PC1 and PC2). (B) Heatmap showing the metabolites that differ significantly between experimental groups (-E – without erythromycin – and +E – with erythromycin) for each strain tested (*Lp*^FlyG2_1.8^ and *Lp*^NIZO2877^) (two-sided t tests, p < 0.05). The compounds are ordered by the metabolite class given by the left scale. Metabolites whose pattern is conserved between strains are framed together in the upper group. (C) Mean ratio fold-change (plotted on a Log2 scale) of metabolites with statistically significant differences (y axis) between experimental conditions (see legend; *: p<0,001). Fold changes and p-values are provided in Supplementary Table 3.

## Discussion

Symbiotic microbes benefit animal health in multiple ways. Different species of gut microbes are known to promote host growth across different animal species in conditions of nutrient scarcity^40,41,44,46^. Bacteria represent a direct energy source (i.e., they serve as protein-rich food)^42,49,62^ and, at the same time, they directly interact with their host as active symbiotic partners^25,43,44,46,62^. As living organisms, host-associated microbes improve digestion of food, provide essential nutrients (e.g., vitamins, amino acids, lipids, etc.), help process the nutritional substrate and boost the host immune response^31,40,41,44,46–48,63,64^. Importantly, different bacterial strains, within the same species, promote host growth to different extents, and this ability directly correlates with the strain-specific ability to thrive in the host nutritional environment^26,49^. Growth promotion is thus among the major beneficial effects mediated by gut microbes. However, the key bacterial features underlying such benefit are currently under debate. What is the predominant factor that drives microbial benefit: being alive or a mere food source? And to what extent strain-specific features impact bacterial potential to promote host development?

In this study, we aimed to create an experimental framework where it is possible to control and test the contribution of the bacterial characteristics primarily involved in promoting animal growth: viability, biomass and strain-specificity. Through the use of erythromycin, a bacteriostatic compound, we studied *L. plantarum*/*Drosophila* association by modulating the physiological state of the bacterial cells. We analysed and ranked the contribution of gut microbes as active partners and as nutritional source, and applied this framework to four different *L. plantarum* strains to test the role of strain-specificity in microbe-mediated benefit. By adding a sub-lethal concentration of erythromycin, we stabilized bacterial growth without affecting the cell nutritional properties and tested the full potential of bacterial biomass to promote larval growth. Our results corroborate the findings of previous studies that the single inoculation of dead (heat-killed or UV-treated) *L. plantarum* strains is not sufficient to recapitulate the beneficial effect exerted by live (and growing) bacteria^43,49^. In addition, we show that different killing treatments (i.e., heat or UV) do not significantly affect the bacterial potential to impact fly development. Nevertheless, being able to assess the potential of live, yet stable, bacterial cells allowed us to reveal strain-specific cell properties involved in promoting larval growth. Specifically, in case of *Lp*^NC8^ and *Lp*^Δdltop^ associations, the single inoculation of stable bacterial cells was sufficient to confer a significantly higher benefit, compared to dead cells, throughout larval development. Contrarily, this was not visible in case of *Lp*^FlyG2.1.8^ and *Lp*^NIZO2877^ associations (Figure 1, Supplementary Figure 2). By daily adding live and dead cells to mimic standard bacterial growth in association with Drosophila, we show that only the recurrent supplementation of viable cells is sufficient to recapitulate the effect of standard bacterial associations, while the addition of dead cells is not (Figure 2, Supplementary Figure 4). These results support what has been previously shown on the contribution of recurrent addition of dead bacterial cells, where daily supply of heat-killed microbes did not alter fly growth or lifespan to the extent of that from standard bacterial association^49,60,62^. Only increasing the quantity of dead bacteria over the amount of live cells was necessary to match or exceed the benefit observed with live microbe inoculation^49,62^. Again, in case of *Lp*^NC8^ and *Lp*^Δdltop^ associations, the supply of dead cells showed a significantly higher benefit compared to axenic conditions, further showing that such strains likely bear cell features and molecules that concur to sustain larval development (Figures 1, 2A). It would be interesting to compare the cell wall structures of these strains with those of *Lp*^FlyG2.1.8^ and *Lp*^NIZO2877^ to identify such components.

Altogether, our results show that bacterial biomass *per se* does not always represent a food source, but the nutritional potential of gut microbes relies on strain-specific cell features. Indeed, we show that strain-specificity contributes to a higher extent to host growth than biomass (Figure 3). In this frame, biochemical properties of bacterial cell walls, as well as structural features, such as cell shape, have been proven to be directly involved in Drosophila growth-promotion. Specifically, D-alanylation of bacterial cell wall teichoic acids (TA), which occurs in *Lp*^NC8^ and not in *Lp*^Δdltop^ strain, is required for promoting Drosophila growth in conditions of nutrient scarcity^45,65^. D-alanylated TA are directly sensed by Drosophila enterocytes triggering intestinal peptidases induction, ultimately concurring to larval growth and maturation. At the same time, the addition of purified cell walls, without the presence of live bacteria, is not sufficient to rescue larval growth to conditions of standard associations^45^. Beyond cell-specific properties, the ability of gut microbes to thrive in the host nutritional environment is known to significantly contribute to microbial benefit. We previously demonstrated that bacterial adaptation to the fly nutritional environment is sufficient to improve *L. plantarum* beneficial effect on larval growth, and that such adaptation leads to faster microbial growth and higher cell counts^26^. Indeed, both strong beneficial strains tested in this study (*Lp*^NC8^ and *Lp*^FlyG2.1.8^) reach higher loads in the fly nutritional environment compared to their isogenic, weakly beneficial strains (*Lp*^Δdltop^ and *Lp*^NIZO2877^)^26,45^ (Figure 2B). Higher loads of beneficial bacteria naturally lead to higher production of beneficial metabolites, which are also needed to sustain animal development^26,41^. However, the presence of such beneficial molecules, without active bacterial cells, barely promotes - and in some conditions even hampers - larval growth^26,60^. By analysing the metabolic profiles of the bacterial strains in different physiological states, we refined the list of bacterial metabolites that are exclusively produced by live and active cells – and not by stable cells – which are likely involved in larval growth promotion. As previously shown, we observed a significant increase in the levels of N-acetyl-amino acids in presence of the strong beneficial strain and *Lp*^FlyG2.1.8^, which explains its higher growth-promoting potential (Figure 4B). At the same time, we revealed further strain-specific differences. We show that the weakly beneficial *Lp*^NIZO2877^ strain consumes more amino acids, sugars and vitamins compared to *Lp*^FlyG2.1.8^, further depriving the host of nutrients required for its growth (Figure 4B, C). This is particularly visible by comparing the levels of galactosamine, N-acetyl-glucosamine and hexose, which resulted to be significantly lower in presence of live and growing *Lp*^NIZO2877^ (Figure 4B). The same trend in digesting nutrients is also visible in presence of *Lp*^FlyG2.1.8^, albeit not significantly different between growing and stable cells (Figure 4B, Supplementary Table 3). Taken together, these results demonstrate that the strain-specific ability to promote animal growth results from a combination of factors, ranging from biochemical and structural cell features, the ability to thrive in the host nutritional environment (i.e., the potential to reach high loads), coupled with the ability to consume less nutrients from the diet, which are then capitalized by the host. It is important to notice that our study is based on association between the host and single bacterial strains. However, animals, including humans, are estimated to harbor hundreds to more than 1,000 bacterial species in their gut intestine^66^, which constantly interact among them and with the host. The metabolic cross-talk that results from their active interplay also largely contributes to the overall benefit exerted by gut microbes^30,31,67^, further highlighting how viability is fundamental to explain gut microbes’ influence inn animal growth.

The symbiotic advantage given by beneficial microbes is multifaceted. In this work, we developed an experimental framework to modulate bacteria physiological states with the aim of clarifying the importance and role of the main bacterial features involved in promoting animal growth. We show that viability is the most important trait contributing to bacteria-mediated benefit in sustaining animal development, over biomass as nutritional source and strain-specific properties, such as growth potential and cell structure. In the future, such framework can be a key tool to modulate bacterial dynamics, viability and growth potential in multiple contexts, such as different diets or within complex communities, by directly targeting species or strains when in association with the host. This will allow to precisely measure the influence of defined species and additional bacterial traits in sustaining animal health.

## Materials and Methods

### Drosophila strains and maintenance

Drosophila *yw* flies were used as the reference strain in this work. Drosophila stocks were cultured at 25°C with 12/12 hour dark/light cycles on a yeast/cornmeal medium containing 50g/l inactivated yeast as described by Storelli et al.^40^. Diet composition: 50g inactivated dried yeast (Bio Springer, Springaline BA95/0-PW), 80g cornmeal (Westhove Farigel, maize H1), 7.2g Agar (VWR#20768.361), 5.2g methylparaben sodium salt (MERCK#106756), 4 mL 99% propionic acid (Sigma-Aldrich P1386) for 1 liter. Standard low-protein diet (LPD) was obtained by reducing the amount of yeast extract to 8g/l. LPD supplemented with erythromycin was prepared by adding 20 μg/ml of erythromycin to standard LPD. Germ-free (GF) stocks were established and maintained as described in Storelli et al.^40^.

### Bacterial strains and culture conditions

The strains used in the present study are listed in Supplementary Table 4. All *L. plantarum* strains were grown in Man, Rogosa, and Sharpe (MRS) medium (broth or supplemented with 15% of microbiological Agar) (Condalab, Spain), overnight at 37°C without shaking. Strains were stored at −80°C in MRS broth containing 20% glycerol.

### MIC tests

Bacteriostatic susceptibility was performed using Liofilchem MIC test strip (Liofilchem S.R.I., Italy) for chloramphenicol, erythromycin, sulfamethoxazole, tetracycline and trimethoprim according to the manufacturers’ instructions. The strips were placed on a Luria Berthani (LB) Agar inoculated with 0.5 ml LB pure colonies’ suspension of *Lp*^NC8^ and Lp^NIZO2877^, separately. Three independent replicates were performed for each antibiotic. Subsequently, the agar plates were incubated at 37°C and observed for Minimum Inhibitory Concentration (MIC) reading after 24 hours. MIC value was determined at the intersection of the strip and the growth inhibition ellipse, as reported by Kowalska-Krochmal et al.^68^.

### Association of *L. plantarum* with *D. melanogaster* larvae

40 embryos collected from GF females were transferred to a fresh LPD GF medium in a 55 mm petri dish. Bacterial strains were cultured to stationary phase (18h) in MRS broth at 37°C. The embryos and the fly medium were mono-associated with 150 μl containing ~1-3 ×10^7^ CFU/ml of the respective bacterial treatment. Emerging larvae were allowed to develop on the contaminated medium at 25°C. For standard mono-association, 150 μl containing ~1-3 ×10^7^ CFU/ml of the respective *L. plantarum* strain resuspended on sterile phosphate buffered saline (PBS, Sigma-Aldrich, USA) were used as inoculum. For the erythromycin treatment (+Ery), 150 μl of ~1-3 ×10^7^ CFUs of the respective *L. plantarum* strain were inoculated on petri dishes containing LPD supplemented with 20 μg/ml of erythromycin. For heat-killed (HK) bacteria treatment, the bacterial pellet obtained from the overnight culture was resuspended in sterile PBS and the bacterial solution was incubated at 60°C for 45min^69^. For inoculation of UV-treated (UVT) bacteria, the bacterial pellet obtained from the overnight culture was resuspended in clear Eppendorf tubes containing sterile PBS and the bacterial solution was exposed to UV light for 20min. After 10min, the solution was mixed thoroughly and re-exposed to the UV light for the remaining 10min.The HK and UVT bacteria solutions were plated out on MRS agar to check for efficient killing prior to inoculation. For the Germ-free treatment (GF), 150 μl of sterile of PBS were inoculated.

### Larval size measurement

Larvae (n ≥ 60) were collected 4 and 7 days after inoculation, washed in distilled water, killed with a short microwave pulse (900W for 15 sec) transferred on a microscopy slide and captured digital images using a Leica Digital Compound Microscope DMD108. The larval longitudinal size (length) was measured using ImageJ software^70^.

### Quantification of bacterial biomass on fly food

10^7^ CFU/ml of *Lp*^NC8^ were inoculated onto 55 mm petri dishes containing LPD fly food, with and without 20 μg/ml of erythromycin and kept for three days (3 independent replicates/condition/time point). 40 mg of fly food were collected at each time point and processed for bacterial DNA extraction using Invisorb® Spin Tissue DNA Mini Kit (Invitec). DNA concentration was calculated by measuring the absorbance of 1 μL of extracted DNA at 260/280 nm using NanoDrop 2000 spectrophotometer. Quantitative PCR was performed in a total of 10 μl on a LightCycler 480 thermal cycler (Roche Diagnostic, Mannheim, Germany) using PowerUp™ SYBR™ Green Master Mix (Applied Biosystems, USA), bacterial DNA and *L. plantarum* specific primer sets designed on *ackA* gene (ackA_F: TAAGACGCAAGATACCCGTG, ackA_R: ACGCACAATCATCAGCTCTT)^26^. The reaction mixture consisted of 0.25 μl of each primer (10 μM each), 5 μl of SYBR GreenER mix, 2 μl of water and 2,5 μl of template cDNA. Each sample has been tested in triplicate. The PCR conditions included 1 cycle of initial denaturation at 95°C for 2 min, followed by 45 cycles of 95°C for 10 sec and 60°C for 60 sec. Absolute quantification of bacterial DNA was conducted as follows: five 1:2 serial dilutions of the standard sample (2 ng/μl of DNA extracted from *L. plantarum*^NIZO2877^ culture) were quantified by Real-time PCR using *ackA* primers (Martino et al., 2018). Each dilution has been tested in triplicate. Melting curves of the detected amplicons were analysed to ensure specific and unique amplification. Standard curves were generated plotting threshold cycle (Ct) values against the log of the standard sample amount. Based on the data obtained from the standard curve, the Ct values of the experimental samples have been used to obtain the DNA concentration (ng/μl) at each time point.

### Recapitulation experiments

To artificially recapitulate the exponential growth of *L. plantarum* on LPD in association with *Drosophila* larvae, we performed daily additions of the respective bacterial strain in the different experimental conditions. First, standard growth of the four *L. plantarum* strains was monitored on LPD containing Drosophila embryos with and without supplementation of 20 μg/ml of erythromycin over the course of 7 days (Supplementary Figure 1). The amount of bacterial cells to inoculate every day was calculated by subtracting the mean CFU/ml in antibiotic conditions from standard conditions (LPD without erythromycin).

Addition of live and non-dividing bacteria (+Ery – daily addition): on day 0, 150 μl containing ~1-3 ×10^7^ CFU/ml of the respective bacterial strain were inoculated on LPD supplemented with 20 μg/ml of erythromycin. Starting from day 3 after mono-association until day 6, 150 μl containing the required load of live bacteria were inoculated daily to recapitulate the exponential growth of each strain in standard conditions. For HT bacteria: on day 0, 150 μl containing ~1-3 ×10^7^ CFU/ml of the respective heat-treated bacterial strain were inoculated on standard LPD with and without 20 μg/ml of erythromycin (Dead + Ery, Dead -Ery, respectively). Starting from day 3 after mono-association until day 6, 150 μl containing the required load of heat-treated bacteria were inoculated daily to recapitulate the exponential growth of each strain in standard conditions.

### Metabolite Profiling

Microtubes containing axenic poor nutrient diet (with and without 20 μg/ml of erythromycin) were inoculated with bacterial suspension (10^3^ CFU/ml). In addition, microtubes containing axenic poor nutrient diet without erythromycin were inoculated with PBS to analyse the metabolite profile of the diet alone. All microtubes have been incubated for 3 days at 25°C. Microtubes were then snap-frozen in liquid nitrogen and stored at −80°C before sending to MS-Omics (https://www.msomics.com/). Five biological replicates per condition were generated. Samples were then extracted and prepared for analysis using MS-Omics’ standard solvent extraction method. The analysis was carried out using a UPLC system (Vanquish, Thermo Fisher Scientific) coupled with a high-resolution quadrupole-orbitrap mass spectrometer (Q Exactive™ HF Hybrid Quadrupole-Orbitrap, Thermo Fisher Scientific). An electrospray ionization interface was used as inoziation source. Analysis was performed in negative and positive ionization mode. The UPLC was performed using a slightly modified version of the protocol described by Doneanu et al.^71^. Data was processed using Compound Discoverer 3.2. (Thermo Fisher Scientific) and Skyline^72^. Identification of compounds were performed at four levels; Level 1: identification by retention times (compared against in-house authentic standards), accurate mass (with an accepted deviation of 3ppm), and MS/MS spectra, Level 2a: identification by retention times (compared against in-house authentic standards), accurate mass (with an accepted deviation of 3ppm). Level 2b: identification by accurate mass (with an accepted deviation of 3ppm), and MS/MS spectra, Level 3: identification by accurate mass alone (with an accepted deviation of 3ppm).

### Statistical analyses

Data representation and analysis was performed using Graphpad PRISM 9 software (www.graphpad.com). A total of 3 to 5 biological replicates were used for all experiments performed in this study in order to ensure representativeness and statistical significance. Two-sided Mann Whitney’s test was applied to perform pairwise statistical analyses between conditions. One-way non-parametric ANOVA (Kruskal-Wallis test) was applied to perform statistical analyses between multiple (n>2) conditions. The PCA model was performed using the descriptive power (DP) of each compound. The DP was calculated as the ratio between the standard deviation within experimental samples and the standard deviation within the quality control (QC) samples. Variables with a ratio higher than 2.5 are most likely to describe variation related to experimental design. Random Forest analysis has been computed on R software v.4.1.0. using *rforest* function in radiant package v.1.4.4, while variable importance has been extracted using *importance* function in ranger package v.0.14.1. The percentage of variable importance on larval size, has been obtained scaling and converting the importance value in percentage using the following commands: *scales∷percent(importance / sum(importance))*.

## Acknowledgments

Research in MEM lab was supported by the STARS@UNIPD Funding Programme (Starting Grant 2017 D.R. 872/2017) of the University of Padova.

## Author contributions

N.S. and M.E.M designed the project; N.S. and S.Y. conducted the experiments; A.Q. conducted the bioinformatics analyses; N.S. and M.E.M analyzed the data and wrote the paper.

## Supplementary files Legends

**Supplementary Figure 1**. (A) *L. plantarum*^NC8^ loads (CFU/ml) monitored over the course of six days on standard fly food (low protein diet, LPD) (-Ery) and on fly food supplemented with 20 μg/ml of erythromycin (+Ery), with (+Fly) and without (-Fly) Drosophila larvae. The result of the non-parametric analysis of covariance (*sm.ancova* function in R) between curves is reported (*p < 0.05). (B) Concentration of *L. plantarum*^NC8^ DNA biomass on standard fly food (low protein diet) (-Ery) and on fly food supplemented with 20 μg/ml of erythromycin (+Ery), quantified by qPCR. Center values in the graph represent means and error bars represent SD.

**Supplementary Figure 2**. (A) Larval longitudinal length measured 4 days after inoculation with 1) live bacterial strains on LPD without erythromycin (Live and growing −Ery); 2) live bacterial strains on LPD supplemented with 20μg/ml of erythromycin (Live and stable +Ery); 2) heat-killed bacterial strains on LPD without erythromycin (Dead – HK); 3) UV-treated bacterial strains on LPD without erythromycin (Dead – UV); and 4) PBS on LPD without erythromycin (GF condition). Asterisks illustrate statistically significant difference on pairwise intra-strain comparisons between standard mono-association (Live and growing −Ery) and the respective treatments (including GF) (***: p<0,0001). Center values in the graph represent means and error bars represent SD. (B) Larval longitudinal length measured 7 days after inoculation with PBS on LPD with erythromycin (GF + Ery) and without erythromycin (GF -Ery) (ns: not significant). Center values in the graph represent means and error bars represent SD.

**Supplementary Figure 3**. (A) Growth trend of the four *L. plantarum* strains monitored over the course of a standard mono-association on LPD. (B) CFUs of the 4 *L. plantarum* strains monitored at day 7 after mono-association on LPD in standard conditions (-Ery), on LPD supplemented with 20 μg/ml of erythromycin with daily addition of live bacterial cells (+ Ery [daily addition]) and on LPD supplemented with 20 μg/ml of erythromycin without daily addition of live bacterial cells (+ Ery [no addition]). Asterisks illustrate statistically significant difference on pairwise intra-strain comparisons between standard mono-association (−Ery) and the respective treatments (***: p<0,0001, ns: not significant). Center values in the graph represent means.

**Supplementary Figure 4**. Larval longitudinal length measured 4 days after inoculation with 1) live bacterial strains on LPD without erythromycin (Live and growing −Ery); 2) daily addition of live bacterial strains on LPD supplemented with 20μg/ml of erythromycin (Live and stable +Ery); 3) daily addition of heat-killed bacterial strains on LPD without erythromycin (Dead -Ery); 4) daily addition of heat-killed bacterial strains on LPD supplemented with 20μg/ml of erythromycin (Dead +Ery) and 5) PBS on LPD without erythromycin (GF condition). Different letters indicate statistically significant differences at p < 0.05. Center values in the graph represent means and error bars represent SD.

**Supplementary Figure 5**. Variable importance obtained through random forest analysis for each bacterial feature in promoting larval growth at day 7 after *L. plantarum*^FlyG2.1.8^ (A), *L. plantarum*^NIZO2877^ (B), *L. plantarum*^NC8^ (C) and *L. plantarum*^Δdltop^ (D) association.

**Supplementary Table 1**. Minimum inhibitory concentration (MIC) range and results for antibiotics used to test susceptibility of *L. plantarum*^NIZO2877^ *L. plantarum*^NC8^ strains.

**Supplementary Table 2**. Multiple comparison results (One-way non-parametric ANOVA Kruskal-Wallis test) obtained by comparing the length of larvae in germ-free (GF) conditions with larvae associated with the respective condition 7 days after mono-association. Results related to Figure 1.

**Supplementary Table 3**. Metabolomic dataset of Drosophila diet inoculated with Lp^NIZO2877^ and Lp^FlyG2.1.8^ separately, with (+E) and without (-E) erythromycin, or with PBS (GF −E). The differences between conditions were analyzed using two-sided t-test. A Benjamini-Hochberg critical value, (i/m)Q, was calculated for each compound, where i = the individual p-value’s rank, m = total number of tests, Q = the false discovery rate. All p-values that are smaller than (i/m)Q are significant.

**Supplementary Table 4**. List of the *L. plantarum* strains used in this study.

## Notes

### Competing Interest Statement

The authors have declared no competing interest.

