## Supplementary figures and images for "Gut microbes predominantly act as symbiotic partners rather than raw nutrients"

### Supplementary Figure 1

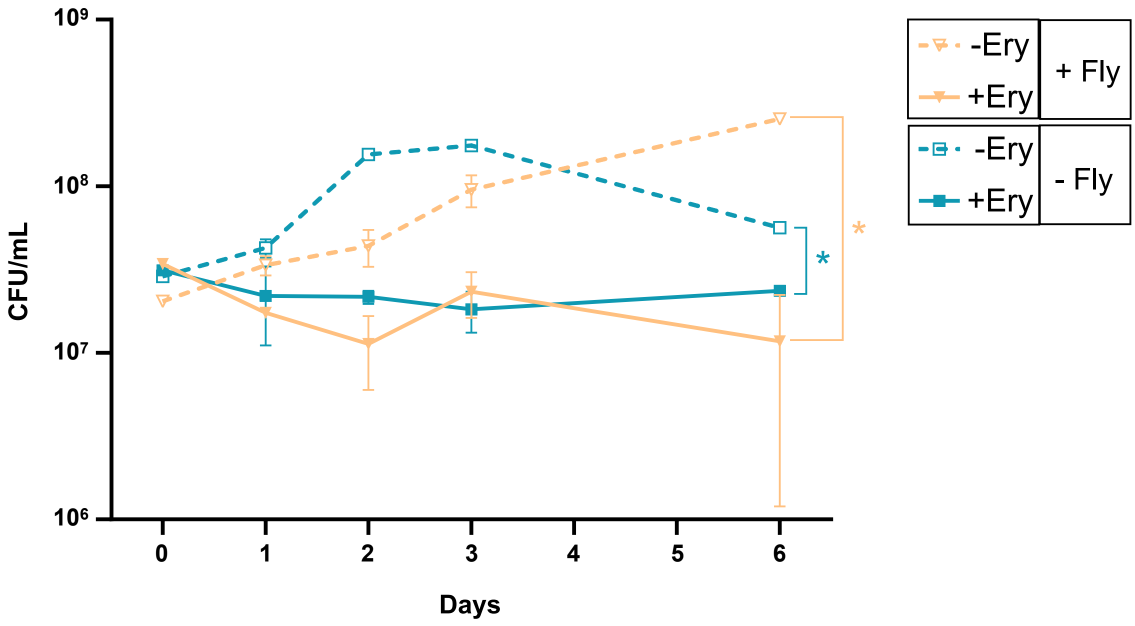

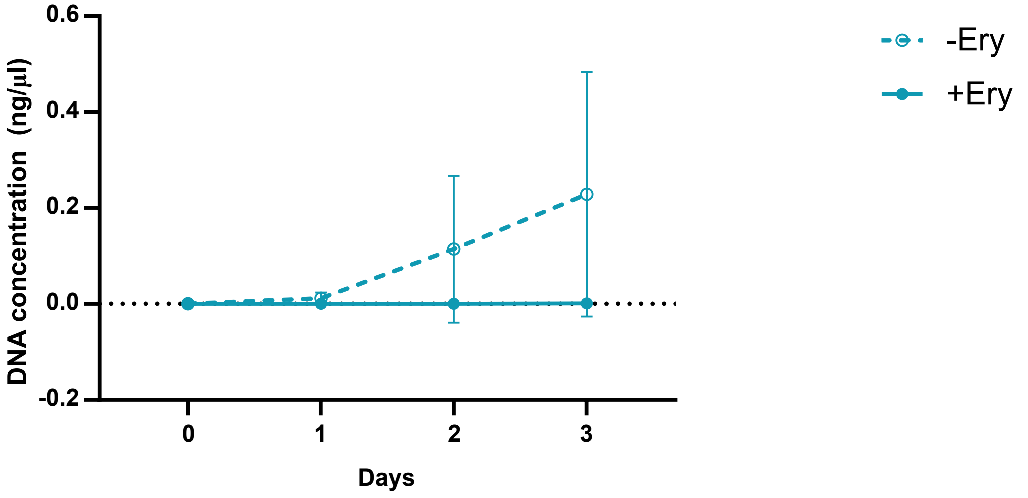

### Supplementary Figure 2

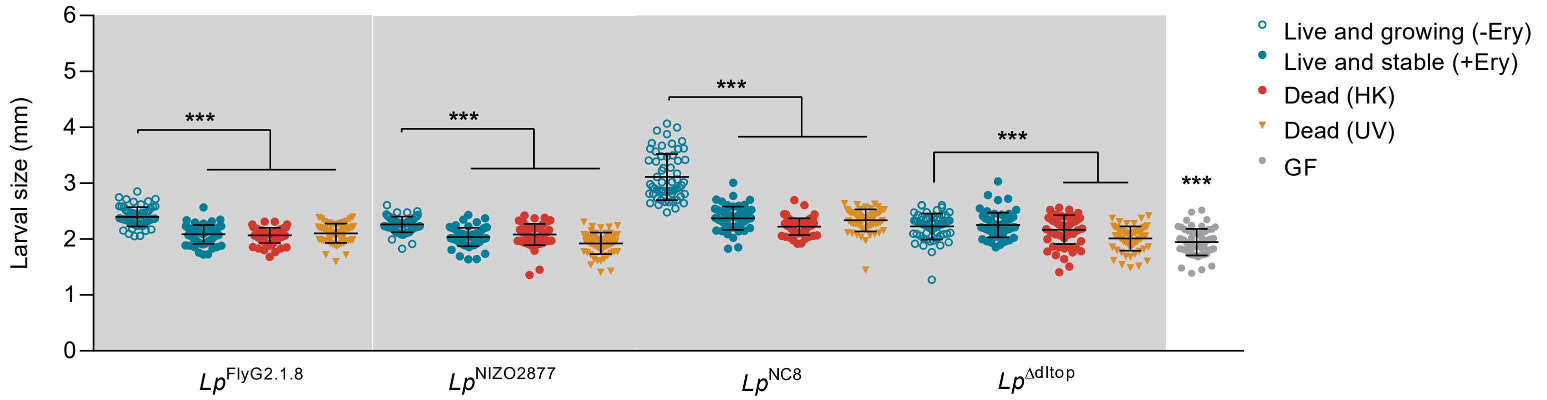

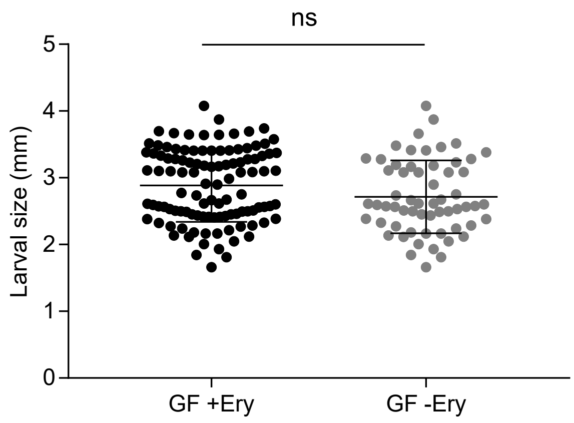

### Supplementary Figure 3

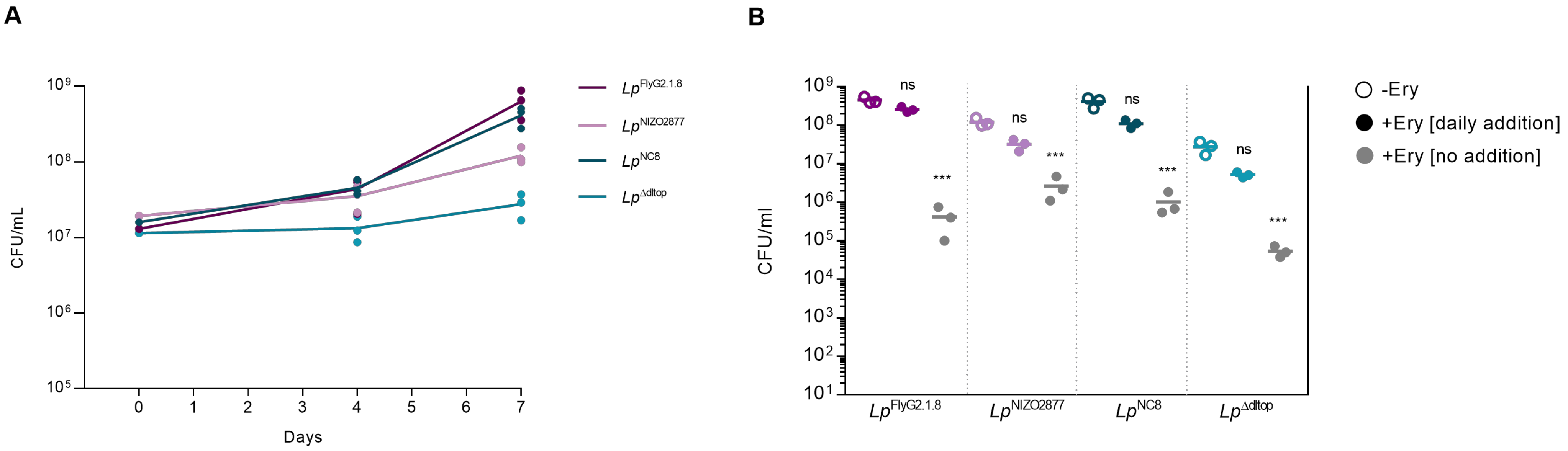

### Supplementary Figure 4

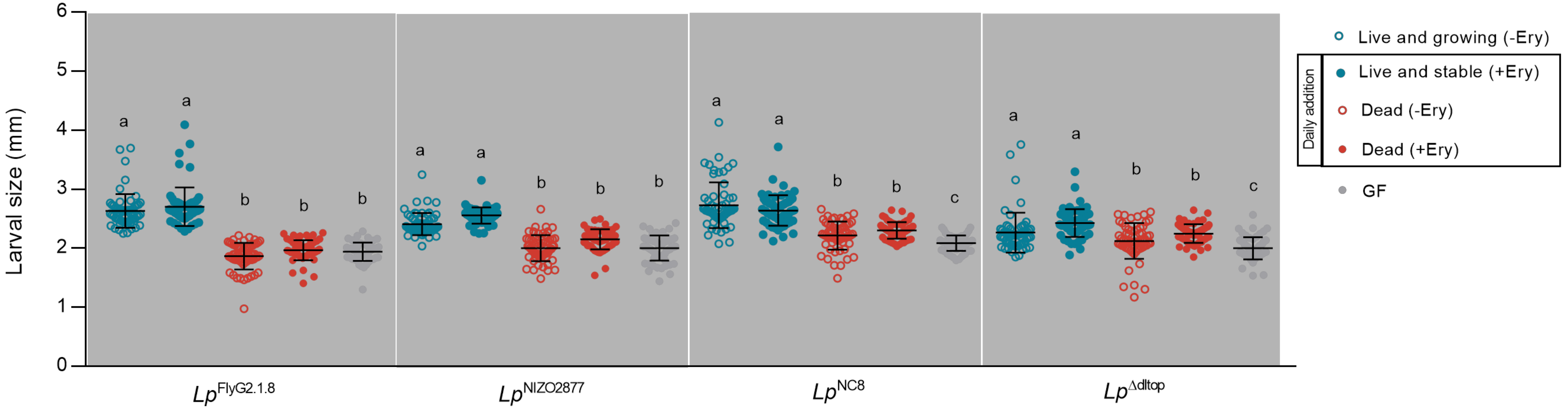

### Supplementary Figure 5

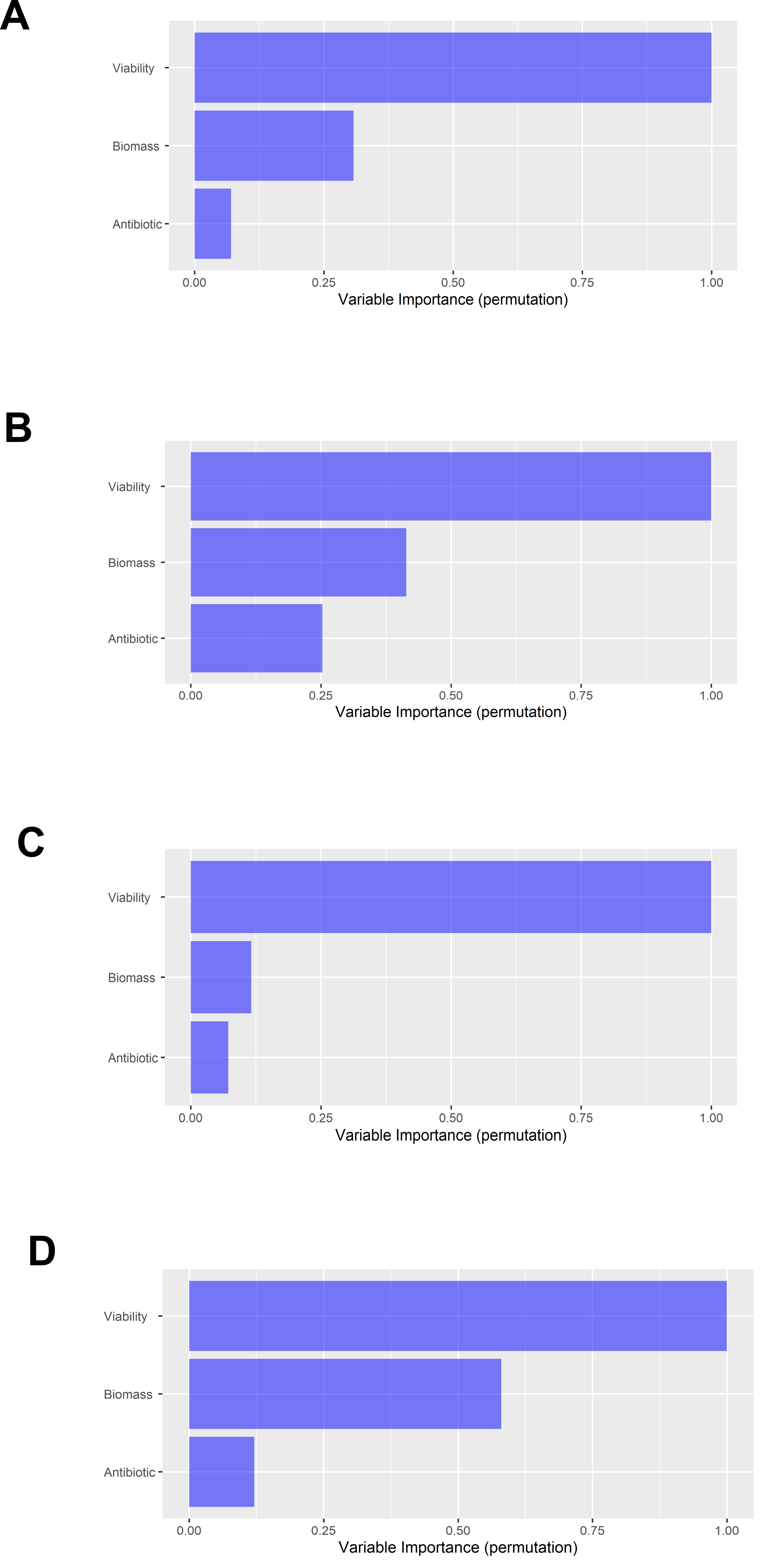
