## Supplementary Table 1 for "Gut microbes predominantly act as symbiotic partners rather than raw nutrients"

| **Antibiotic** | **Code** | **MIC range tested (μg/ml)** | ***L. plantarum*^NIZO2877^ MIC (μg/ml)** | ***L. plantarum*^NC8^ MIC (μg/ml)** |
| --- | --- | --- | --- | --- |
| Chloramphenicol | C | 0.016 - 256 | 4 | 2 |
| Erythromycin | E | 0.016 - 256 | 0.25 | 0.25 |
| Sulfamethoxadole | SMX | 0.064 - 1024 | Resistant | Resistant |
| Tetracycline | TE | 0.016 - 256 | 2 | 2 |
| Trimethoprim | TM | 0.002 - 32 | 0.38 | 0.5 |
