## Supplementary Table 4 for "Gut microbes predominantly act as symbiotic partners rather than raw nutrients"

| **Strain** | **Description** | **Reference** |
| --- | --- | --- |
| *L. plantarum*^NC8^ | - Isolated from grass silage  - Strong growth-promoting strain |  |
| *L. plantarum*^Δdltop^ | - Derived from *L. plantarum*^NC8^ through transposon mutagenesis  - Isogenic to *L. plantarum*^NC8^  - Weak growth-promoting strain |  |
| *L. plantarum*^NIZO2877^ | - Isolated from Vietnamese hotdog  - Weak growth-promoting strain |  |
| *L. plantarum*^FlyG2.1.8^ | - Experimentally evolved from *L. plantarum*^NIZO2877^  - Isogenic to *L. plantarum*^NIZO2877^  - Strong growth-promoting strain |  |
